# Distinct states of proinsulin misfolding in MIDY

**DOI:** 10.1101/2021.05.10.442447

**Authors:** Leena Haataja, Anoop Arunagiri, Anis Hassan, Kaitlin Regan, Billy Tsai, Balamurugan Dhayalan, Michael A. Weiss, Ming Liu, Peter Arvan

**Author notes:** To whom correspondence may be addressed: Peter Arvan MD PhD, Division of Metabolism, Endocrinology & Diabetes, University of Michigan, Brehm Tower rm 5112, 1000 Wall St. Ann Arbor, MI 48105, email.

## Abstract

A precondition for efficient proinsulin export from the endoplasmic reticulum (ER) is that proinsulin meets ER quality control folding requirements, including formation of the Cys(B19)-Cys(A20) “interchain” disulfide bond, facilitating formation of the Cys(B7)-Cys(A7) bridge. The third proinsulin disulfide, Cys(A6)-Cys(A11), is not required for anterograde trafficking, i.e., a “lose-A6/A11” mutant [Cys(A6), Cys(A11) both converted to Ser] is well secreted. Nevertheless, an unpaired Cys(A11) can participate in disulfide mispairings, causing ER retention of proinsulin. Among the many missense mutations causing the syndrome of Mutant *INS* gene-induced Diabetes of Youth (MIDY), all seem to exhibit perturbed proinsulin disulfide bond formation. Here we have examined a series of seven MIDY mutants [including G(B8)V, Y(B26)C, L(A16)P, H(B5)D, V(B18)A, R(Cpep+2)C, E(A4)K], six of which are essentially completely blocked in export from the ER in pancreatic β-cells. Three of these mutants, however, must disrupt the Cys(A6)-Cys(A11) pairing to expose a critical unpaired cysteine thiol perturbation of proinsulin folding and ER export, because when introduced into the proinsulin lose-A6/A11 background, these mutants exhibit native-like disulfide bonding and improved trafficking. This maneuver also ameliorates dominant-negative blockade of export of co-expressed wild-type proinsulin. A growing molecular understanding of proinsulin misfolding may permit allele-specific pharmacological targeting for some MIDY mutants.

## Introduction

Patients bearing autosomal dominant diabetogenic mutations in the *INS* gene [1, 2] develop a syndrome referred to as Mutant *INS* gene-induced Diabetes of Youth [MIDY [3–5]; alternative designations include non-autoimmune type 1 diabetes, MODY10 [6, 7] (OMIM #613370), as well as other names [8]]. MIDY mutations cause proinsulin misfolding in the endoplasmic reticulum (ER) [1, 9]; moreover, MIDY mutants directly associate with wild-type (WT) “bystander” proinsulin, thereby imposing impaired ER exit onto those molecules as well [10–15].

Pancreatic β-cells can synthesize > 6000 new proinsulin molecules/second [16]. To achieve insulin bioactivity, proinsulin must form three native disulfide bonds: Cys(B7)-Cys(A7), Cys(B19)-Cys(A20), and Cys(A6)-Cys(A11) [17] (Fig. 1); thus the β-cell ER supports formation ≥ 18,000 disulfide bonds/second, designed to accommodate physiological levels of insulin production [5]. However, whereas three native disulfide bonds are required for the ultimate bioactivity of insulin, only two of these bonds are needed for proinsulin export from the ER [18]. Indeed, we developed a proinsulin mutant known as “lose-A6/A11” which rapidly forms the two native disulfide bonds [Cys(B7)-Cys(A7) and Cys(B19)-Cys(A20)], undergoes intracellular transport, and localizes to secretory granules in β-cells — this variant is secreted with an efficiency comparable to that of WT proinsulin [19]. On the other hand, we also recognize that MIDY mutant mice bearing the heterozygous *Ins2*-*Munich* allele — encoding proinsulin-C(A6)S — develop severe insulin-deficient diabetes [20], indicating that an unpaired proinsulin-Cys(A11) creates an impediment to proinsulin trafficking through disulfide mispairings [21] that can result in catastrophic proinsulin misfolding [19].

**Figure 1.**
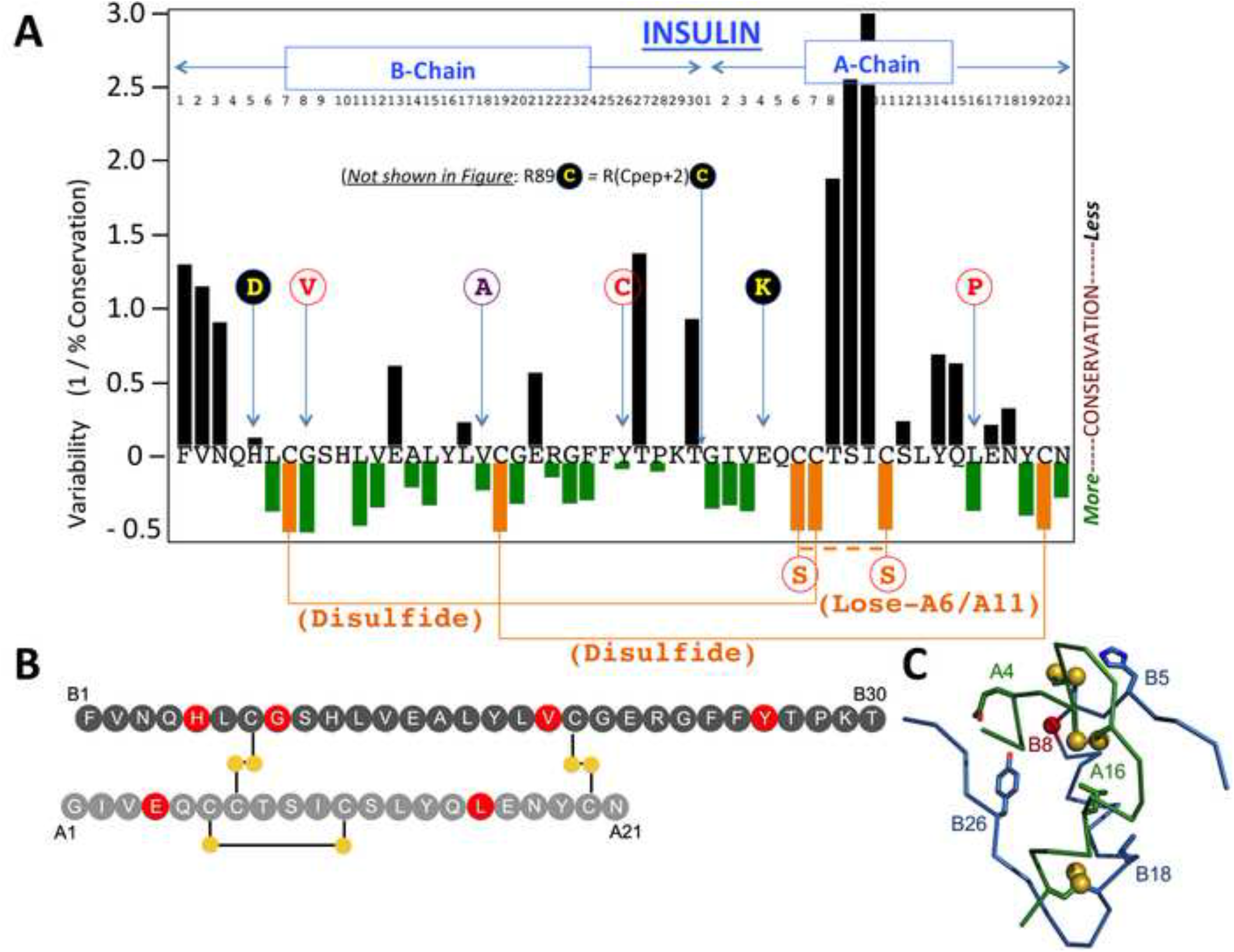
Position of missense MIDY mutants studied in this manuscript, shown relative to the peptide sequence of human insulin. **A)** Numbering of residues of the insulin B-chain, and insulin A-chain (referring to peptide sequence as residues B1 to B30, and A1 to A21, respectively) is shown near the top of the Figure (*the C-peptide is not shown*). The X-axis indicates the primary structure of mature human insulin (single-letter code) organized as a map of the degree of sequence variability at each position, from [28] — negative values indicate higher conservation with increasingly positive values showing greater levels of variability. Cys residues are shown in gold and native disulfide bonds indicated as solid gold lines. Mutations altering endogenous human proinsulin residues, engineered in this study, are shown as circles in single-letter code. The double-mutant C(A6)S,C(A11)S is referred to in this manuscript as ‘lose-A6/A11’, and a dashed gold line indicates what would normally be a native disulfide bond that cannot be made in this double-mutant. Replacement of Val(B18) with Ala is shown in a unique color (purple) designating its unique behavior described herein. G(B8)V, Y(B26)C, and L(A16)P are each shown in red font within white circles, signifying resistance to secretion rescue in the lose-A6/A11 background. The remaining studied mutations are shown as yellow font in black circles, signify MIDY mutants that exhibit significant secretion rescue within the lose-A6/A11 background. A dibasic cleavage site separates the C-peptide from the A-chain, the second residue of which is the site of the R(Cpep+2)C MIDY mutation, which has also been described as R(89)C. **B)** Two-chain structure of insulin with B-chain (top; dark gray) and A-chain (bottom; light gray). Sites of mutation within the insulin moiety of proinsulin considered in this study are highlighted in red. Disulfide bridges are shown as black lines with sulfur atoms as filled gold circles. **C)** Crystallographic T-state protomer (Protein Databank entry 4INS, 2-Zn molecule-1) with B-chain in blue, A-chain in red, sulfur atoms engaged in three cystines (B7-A7, B19-A20 and A6-A11) shown in gold, and sites of mutation within the insulin moiety of proinsulin considered in this study denoted (the C_a_ carbon of Gly^B8^ is shown as a red ball).

Distinct MIDY mutant alleles are associated with diabetes onset ranging from neonatal life through adulthood [2, 8]. In the current study, we sought to determine whether Cys residues from the A6/A11 tandem might contribute to proinsuiln failure in MIDY. Here, we provide evidence for two main subsets of misfolded MIDY mutants that we propose are distinguished by their competence in forming the Cys(A6)-Cys(A11) disulfide. In one subset, impaired closure of the Cys(A6)-Cys(A11) bridge leads directly to aberrant intermolecular disulfide-linked complexes, blocking proinsulin export from the ER and dominantly retaining co-expressed WT proinsulin.

## METHODS

### Materials, Antibodies, and Human Insulin Immunoassay

See Resources Table. We make special note that the manufacturer indicates that the human insulin STELLUX ELISA assay (ALPCO 80-INSHU-CH01) has a cross-reactivity with human insulin of 100%; human proinsulin undetectable; mouse insulin undetectable; rat insulin-1 0.49%, and rat insulin-2 undetectable.

### Plasmids encoding mutant proinsulins

Plasmids encoding myc-tagged human proinsulin, myc-tagged human “lose-A6/A11” proinsulin and hPro-CpepSfGFP (Resources Table) were used as templates for mutagenesis using the QuikChange site-directed mutagenesis kit, with resulting plasmids confirmed by direct DNA sequencing.

### Cell Culture and transfection: 293T, INS832/13, and Min6 cells

INS832/13 cells (see Key Resources Table) were cultured in RPMI-1640 medium supplemented with 10% FBS, 1 mM sodium pyruvate, 0.05 mM β-mercaptoethanol, 10 mM HEPES pH 7.35, and penicillin/streptomycin. Min6 cells (see Key Resources Table) were cultured in DMEM (25 mM glucose) supplemented with 10% FBS, 1 mM sodium pyruvate, 0.1 mM β-mercaptoethanol, and penicillin/streptomycin. 293T cells (see Key Resources Table) were cultured in DMEM supplemented with 10% calf serum and penicillin/streptomycin. Cells were transfected with lipofectamine (Thermo-Fisher Scientific).

### SDS-PAGE and Western blotting

At 24 or 48 h post-transfection, cells were washed with PBS and lysed in Laemmli sample buffer or RIPA buffer (10 mM Tris pH 7.4, 150 mM NaCl, 0.1% SDS, 1% NP40, 2 mM EDTA) including protease/phosphatase inhibitor cocktail (Sigma-Aldrich). Samples were resolved by SDS-PAGE in 4–12% Bis-Tris NuPAGE gels under either nonreducing or reducing conditions. Completed nonreducing gels were incubated in 30 mM dithiothreitol (DTT) for 10 min at room temperature prior to electrotransfer to nitrocellulose. Development of immunoblots used enhanced chemiluminescence, captured with a Fotodyne gel imager, quantified using ImageJ.

### Metabolic labeling, immunoprecipitation, and Tris-tricine-urea-SDS-PAGE

Transfected INS832/13 were incubated in media lacking Cys and Met for 30 min; ^35^S-pulse-labeled for 10 - 30 min at 37°C (Tran^35^S label, Perkin Elmer) and chased in complete growth media; at chase time, media were collected. Before lysis, cells were washed with ice-cold PBS containing 20 mM N-ethyl maleimide (NEM), and then lysed in RIPA buffer (25 mM Tris, pH 7.5, 100 nM NaCl, 1% Triton X-100, 0.2% deoxycholic acid, 0.1% SDS, 10 mM EDTA) [or, in Fig. 7C, 10 mM Tris pH 7.4, 150 mM NaCl, 1% NP-40, 0.1% SDS, 2 mM EDTA] containing 2 mM NEM and a protease inhibitor cocktail. Cell lysates, normalized to trichloroacetic acid-precipitable counts, were pre-cleared with pansorbin and immunoprecipitated with anti-Myc, anti-insulin, or anti-GFP antibodies and protein A-agarose overnight at 4°C. Immunoprecipitates were analyzed by nonreducing / reducing Tris–tricine–urea–SDS-PAGE, or 4-12% gradient Nu-PAGE. Gels were fixed and dried, followed by phosphorimaging or autoradiography; bands were quantified using ImageJ.

### Live cell Microscopy of GFP-tagged proinsulins

Transfected INS832/13 cells were cultured in LabTek-II coverglass chambers (Nunc). At 24 h post-transfection live cell fluorescence was examined using a Nikon A1 Confocal Microscope. Multiple image/fields were captured for each construct; raw images were processed using NIS-Elements Viewer 5.21 software.

### ER stress response as measured by BiP-Luciferase

Cells were co-transfected with plasmids encoding human proinsulin construct, BiP promoter-firefly luciferase reporter, and renilla luciferase, at a DNA ratio of 100:10:1. At 48 h post-transfection, cells were lysed and analyzed for firefly luciferase normalized to renilla luciferase activity (Promega Dual Luciferase assay).

### Statistical Analysis

Results were calculated as mean ± s.d.. Statistical analyses employed two-tailed unpaired Student’s *t*-test (Figs. 3B, 7E), or one-way ANOVA followed by Dunnett’s for multiple comparisons to WT (Figs. 3C, 4B, 4D, 8C, S2, S3), or Tukey’s test for multiple comparisons between all samples (Figs. 6B, 6E, S4), using GraphPad Prism v.8. A *p*-value of < 0.05 was taken as significant.

**Figure 2.**
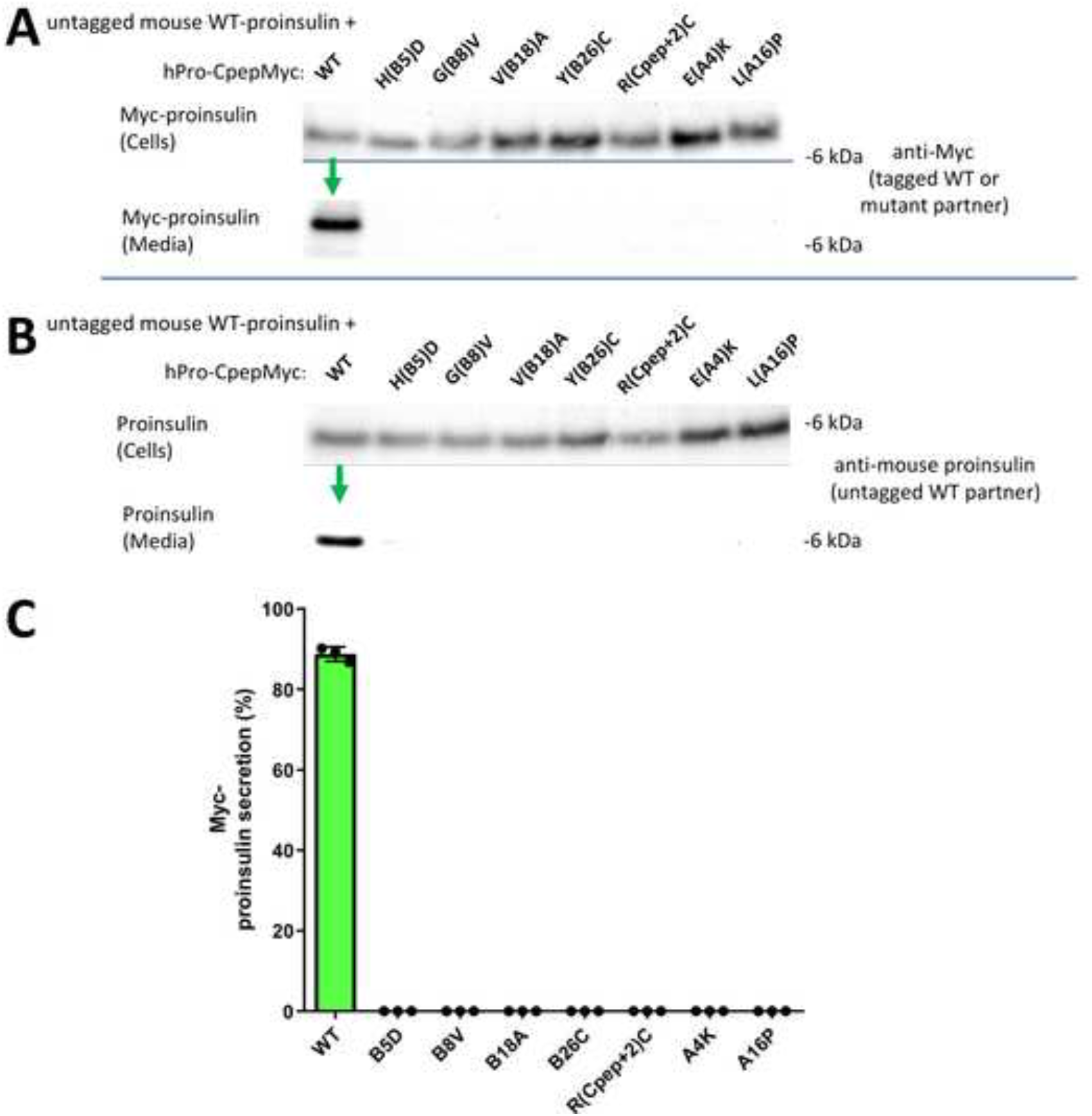
Co-expression of human mutant proinsulin with mouse wild-type proinsulin demonstrates dominant-negative inhibition of proinsulin export. 293T cells were co-transfected to express mouse WT proinsulin and myc-tagged human WT or MIDY mutant proinsulin at a plasmid ratio of 1 : 2, respectively [10]. Beginning at 6 h post-transfection, cell culture media was collected for one day, and the cells were then lysed. Cellular content and secretion of proinsulin was proportionately loaded and resolved by SDS-PAGE. **A)** Cell and media content of myc-tagged proinsulins by immunoblotting with anti-myc antibody; tagged-WT proinsulin was secreted (green arrow). **B)** Cell and media content of co-expressed untagged mouse WT proinsulin by immunoblotting with rodent-specific anti-proinsulin mAb; untagged-WT proinsulin was secreted (green arrow). **C)** Secretion of WT or MIDY mutant human proinsulins were quantified from three independent replicate experiments conducted as in panel A.

**Figure 3.**
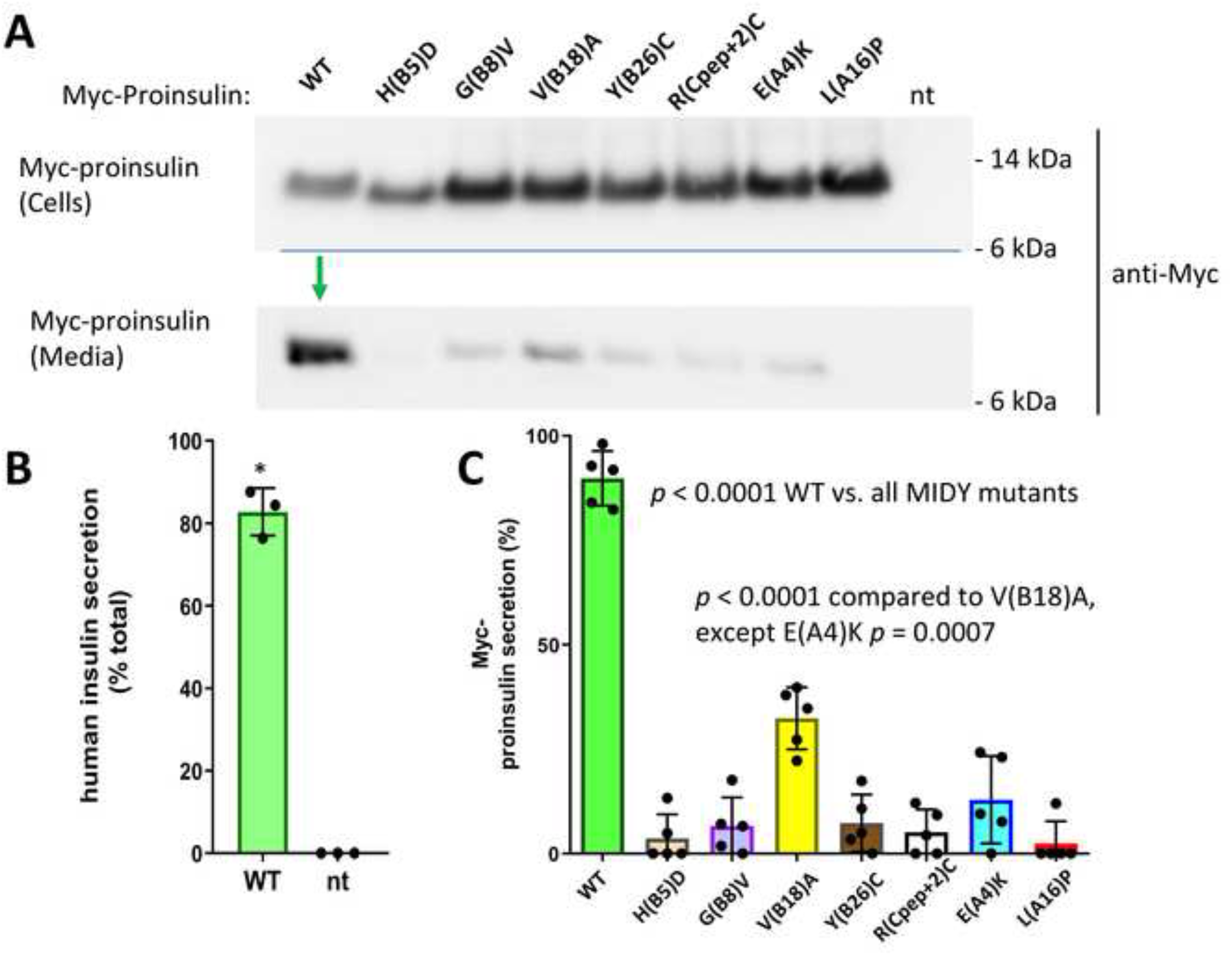
Mutant human proinsulin export in β-cells. Min6 β-cells were transfected with plasmids encoding myc-tagged human WT or MIDY mutant proinsulins, as indicated. At 48 h post-transfection, cell culture media was collected overnight, and the cells were lysed. Cell lysates and secretion media were proportionately loaded and resolved by SDS-PAGE and electrotransfer to nitrocellulose. **A)** Cell and media content of myc-tagged proinsulins by immunoblotting with anti-myc antibody, demonstrating abundant secretion of WT human proinsulin (green arrow). **B)** Confirming that myc-tagged WT proinsulin is competent in the biosynthesis of authentic (human) insulin, secretion was measured by human-specific insulin ELISA; endogenous rodent insulin (from nontransfected cells, ‘nt’) cannot be detected in the assay (mean ± s.d. shown from 3 independent experiments, \**p* < 0.05). **C)** Secretion of myc-tagged human WT and MIDY mutant proinsulins like that shown in panel A were quantified by scanning densitometry (n = 5 independent experiments); with *p*-values on the figure.

**Figure 4.**
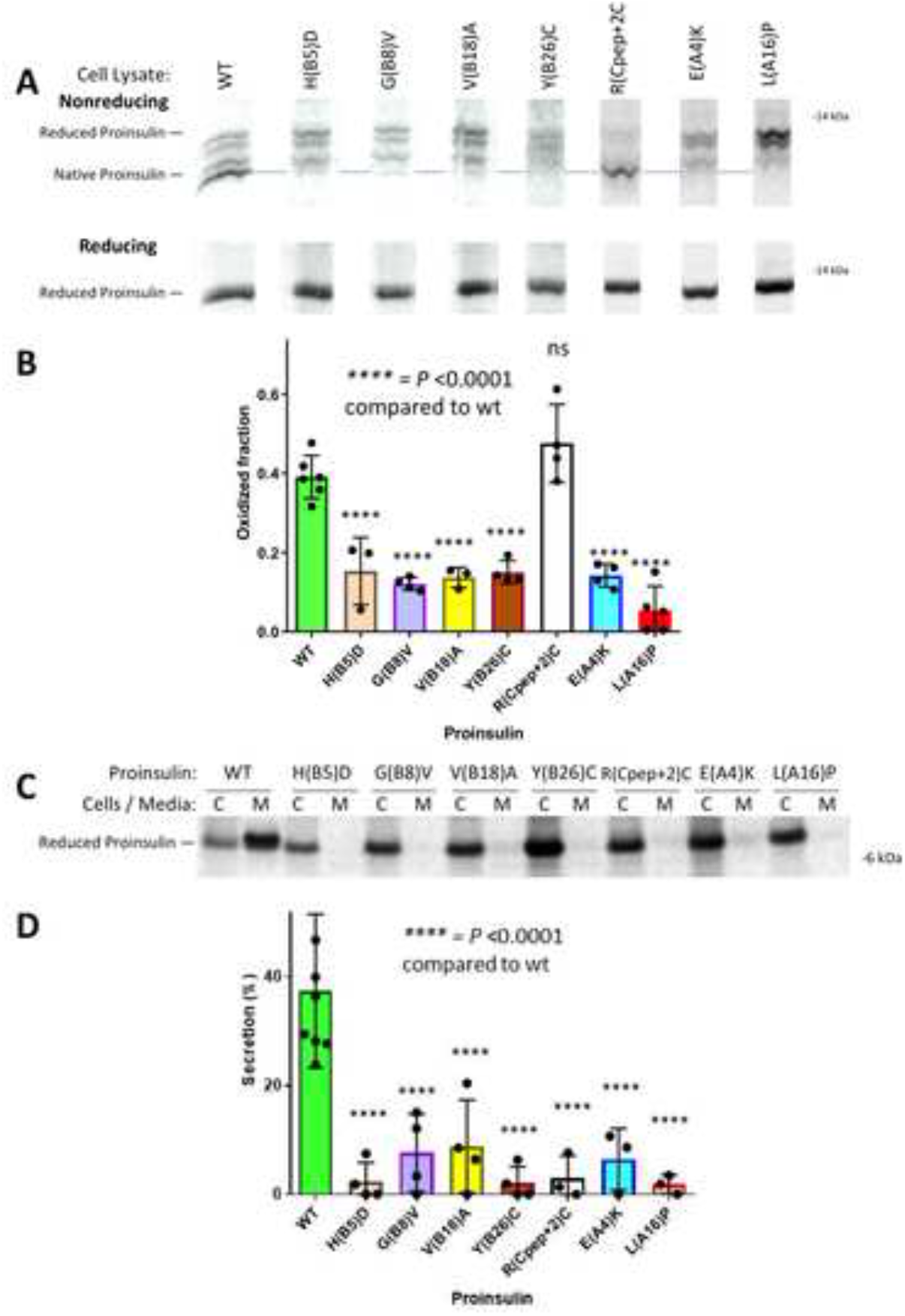
Oxidation and secretion of newly-synthesized mutant proinsulin. INS832/13 β-cells transfected to express myc-tagged human WT or MIDY mutant proinsulins were pulse-labeled with ^35^S-labeled amino acids for 30 min and either lysed immediately (zero chase time, panels A, B) or chased in complete media for 3 h (panels C, D). Chase media were collected and the cells were lysed (see Methods). **A)** Cell lysates were immunoprecipitated with anti-Myc antibodies and analyzed by nonreducing (upper panel) or reducing Tris–tricine–urea–SDS-PAGE (lower panel) and phosphorimaging. A dashed light-blue line shows the position of the oxidized proinsulin monomer; all lanes come from the same gel image. **B)** At the zero chase time, the fastest-migrating form of oxidized proinsulin, as a fraction of total proinsulin, was quantified (mean ± s.d. from at least 3 independent experiments). **C)** After 3h chase, cell lysates (C) and media (M) were immunoprecipitated with anti-Myc-antibodies and analyzed by reducing SDS-PAGE and phosphorimaging. **D)** Experiments like that shown in panel C were performed in at least 3 independent experiments, and secretion quantified (mean ± s.d.); with *p*-values shown on the figure.

**Figure 5.**
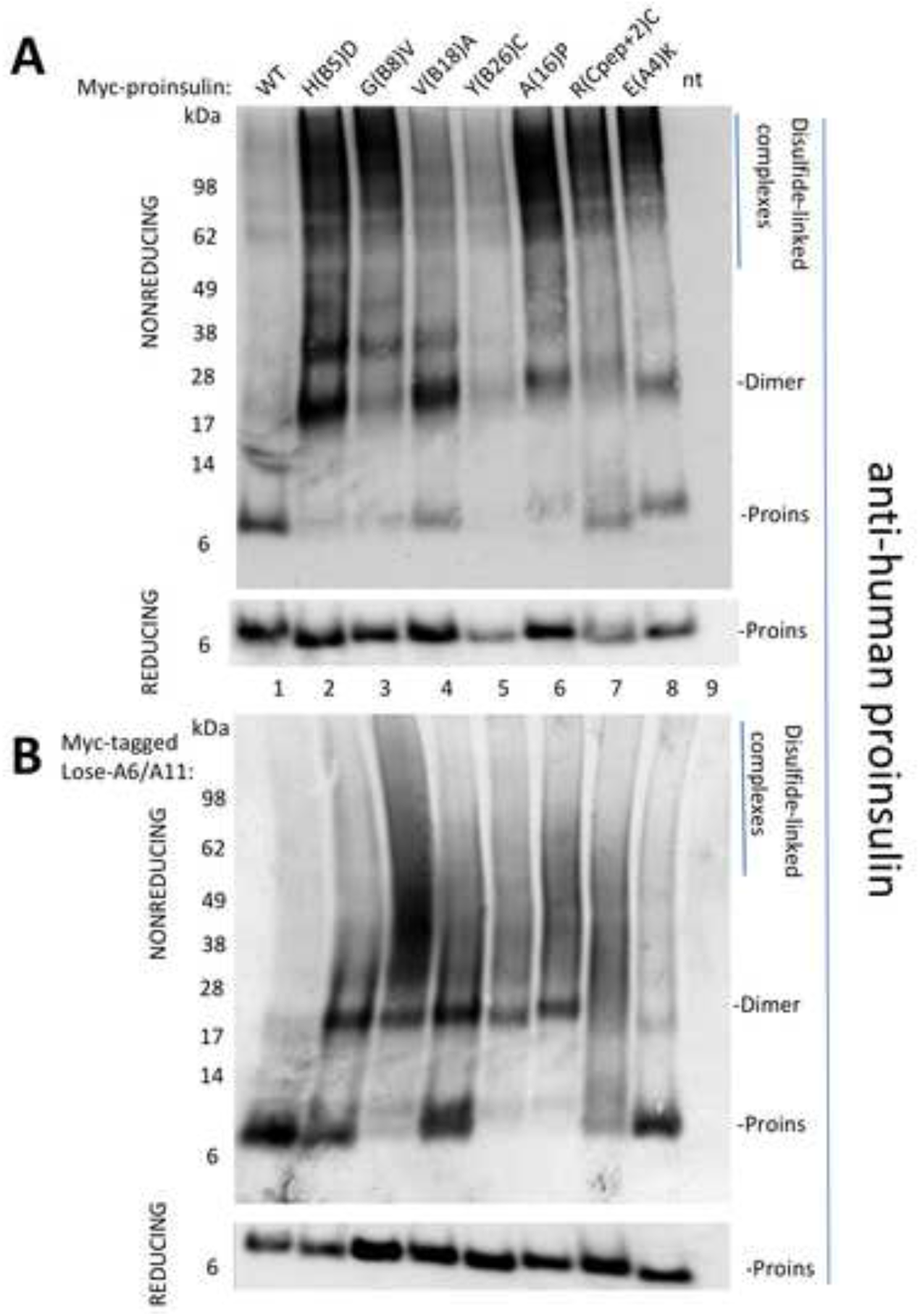
Disulfide-linked complexes of MIDY proinsulins tend to be rescued in the lose-A6/A11 background. **A)** Min6 β-cells were transiently transfected to express myc-tagged human WT or MIDY mutant proinsulins, as indicated. At 48 h post-transfection, the cells were lysed and analyzed by nonreducing or reducing SDS-PAGE (NuPAGE), electrotransfer, and immunoblotting with anti-human-proinsulin mAb (directed against a sequence spanning the human B-chain/C-peptide junction). From nonreducing gels, proinsulin was detected either as monomer (band labeled ‘Proins’) or in a ladder of disulfide-linked dimers, trimers and higher-molecular weight disulfide-linked complexes (as indicated). **B)** Min6 β-cells were transiently transfected to express myc-tagged lose-A6/A11 proinsulins bearing WT or MIDY mutations, and analyzed exactly as in panel A. Note that the antibody does not detect any endogenous proinsulin of nontransfected mouse β-cells under either nonreducing or reducing conditions (lane 9, “nt”).

**Figure 6.**
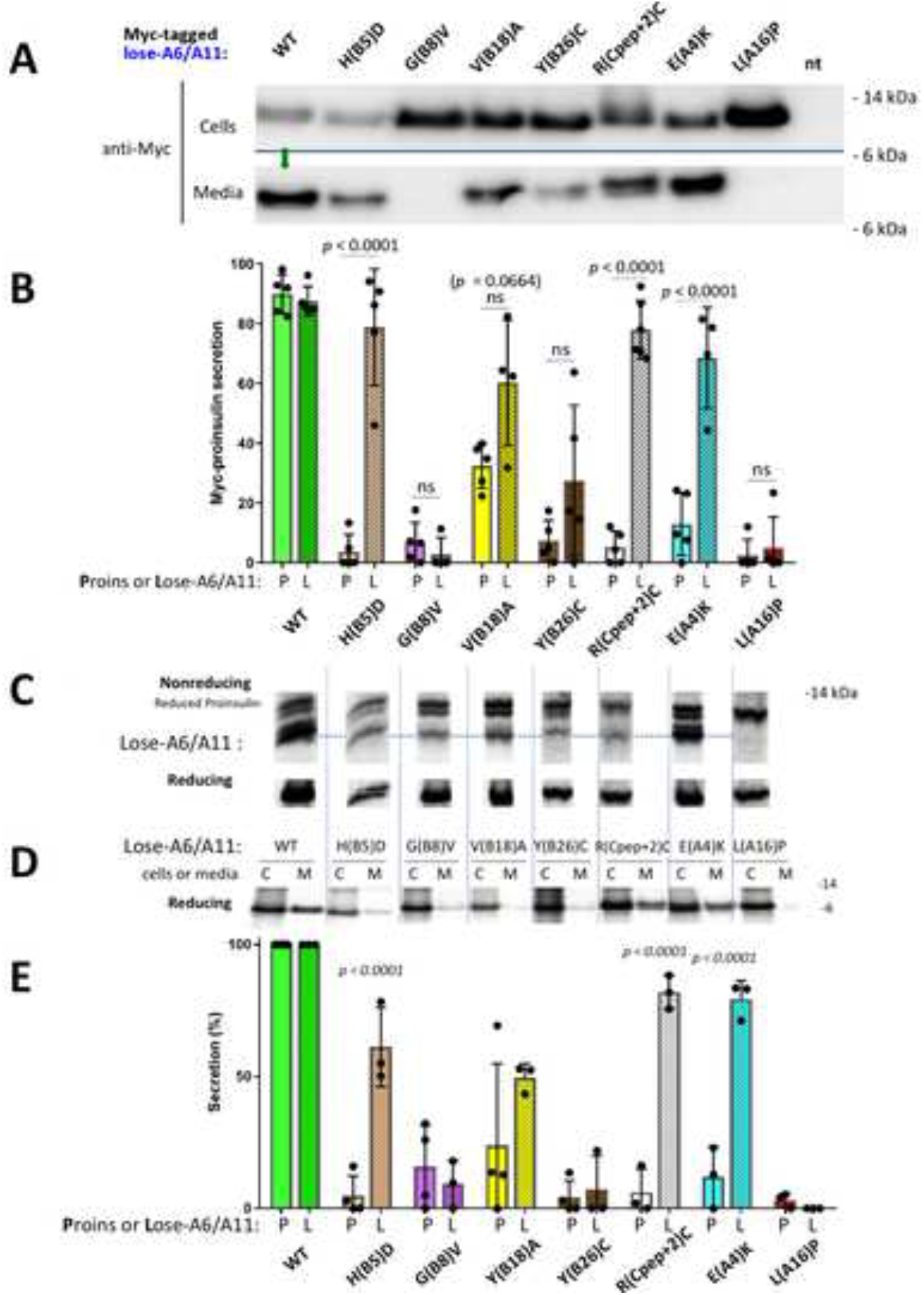
Some mutant proinsulins in the lose-A6/A11 background are secreted. Min6 β-cells were transfected to express myc-tagged human lose-A6/A11 WT proinsulin, or that bearing MIDY mutants as indicated. At 48 h post-transfection, cell culture media was collected overnight, and the cells were lysed. Cell lysates and media were proportionately loaded and resolved by SDS-PAGE. **A)** Cell and media content of myc-tagged proinsulins resolved under reducing conditions, analyzed by immunoblotting with anti-Myc antibody; WT proinsulin is secreted (green arrow). **B)** Secretion of lose-A6/A11 constructs like those in panel A were quantified in comparison to that of the same MIDY mutants in the WT proinsulin background, from Fig 3C. Secretion from at least three independent experiments is shown; *p*-values (on the figure) were analyzed for the lose-A6/A11 background (hatched bars) compared to the proinsulin background (solid bars). **C+D)** INS832/13 cells were transfected to express myc-tagged human lose-A6/A11 proinsulin or that bearing MIDY mutants as indicated. At 48 h post-transfection, the cells were pulse-labeled with ^35^S-labeled amino acids for 30 min. **C)** Samples were lysed immediately, processed as in Fig. 4A, and analyzed by nonreducing (upper panel) or reducing Tris–tricine–urea–SDS-PAGE (lower panel) and phosphorimaging. **D)** Samples were chased for 90 min; cell lysates (C) and media (M) were immunoprecipitated with anti-Myc antibodies and processed as in Fig. 4C (reducing SDS-PAGE and phosphorimaging). **E)** Secretion of the MIDY mutants in the WT proinsulin background (solid bars) or lose-A6/A11 background (hatched bars) was quantified (mean ± s.d. from at least 3 independent experiments). In each experiment, for the insulin chain sequence not bearing a MIDY mutation, secretion was set to 100%.

**Figure 7.**
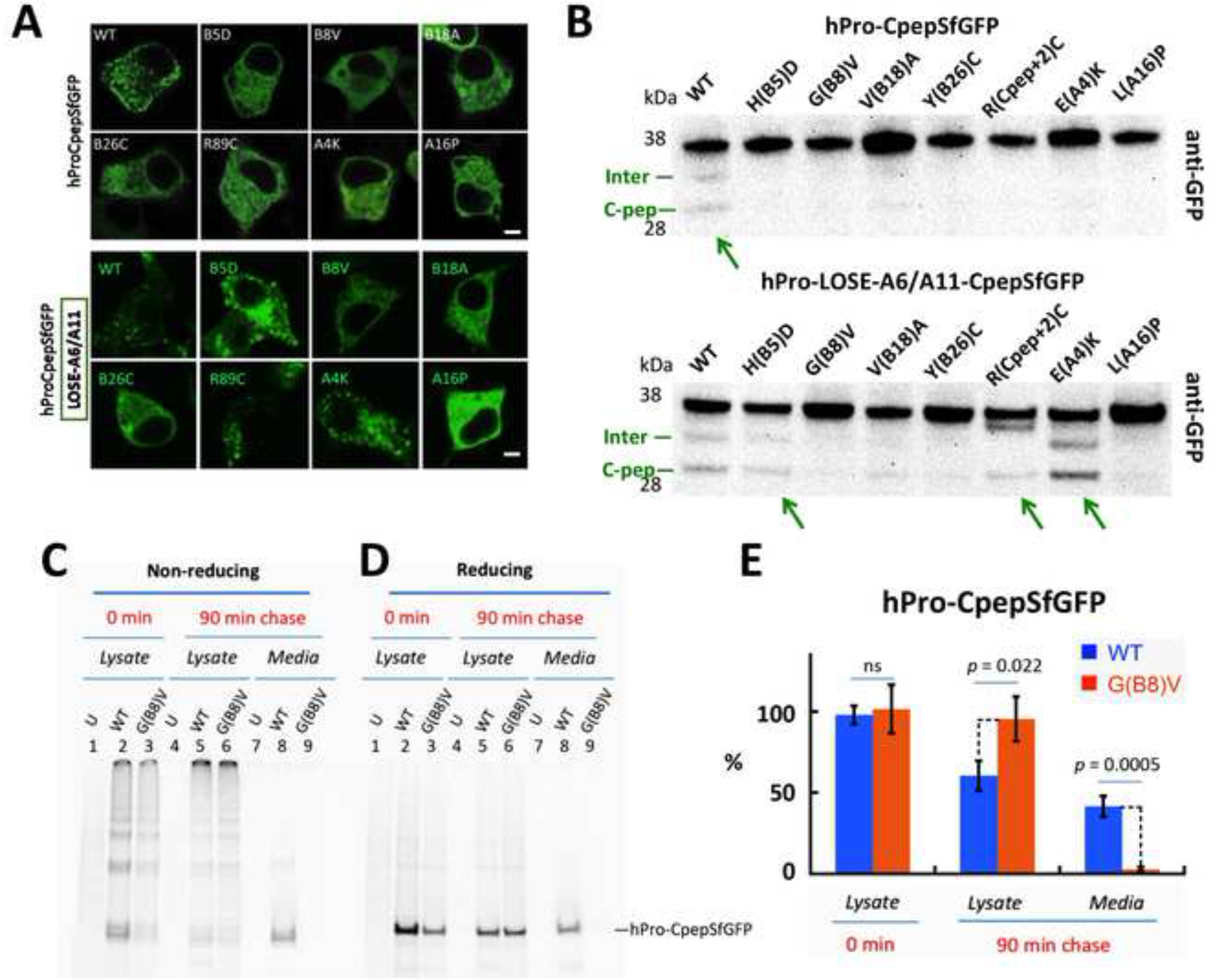
Intracellular distribution of MIDY mutant proinsulins in live β-cells. **A)** Live cell imaging of INS832/13 cells transfected to transiently express fluorescent WT or MIDY proinsulins (and their derived fluorescent protein products) bearing superfolder-GFP embedded within the C-peptide. *Upper panels* show each construct contained within the hPro-CpepSfGFP background. *Lower panels* show the same WT and MIDY mutants in which the hPro-CpepSfGFP bears the lose-A6/A11 substitutions. White scale bar = 10 µm. (Additional “volume views” using 3D-tilt images are shown in Supplemental Fig. S5.) **B)** Anti-GFP immunoblotting of cell lysates from INS832/13 cells transiently transfected as in panel A, and resolved by reducing SDS-PAGE. *Upper panels* show the dark full-length hPro-CpepGFP band for each construct. The construct bearing the WT insulin chains also demonstrates an endoproteolytic processing intermediate (‘Inter’) and the fully-mature CpepSfGFP (green arrow). *Lower panels* show the same constructs in the lose-A6/A11 background, demonstrating more obvious endoproteolytic processing for H(B5)D, R(Cpep+2)C, and E(A4)K (green arrows). A representative experiment (of three) is shown. **C)** 293T cells were either untransfected (‘U’) or transiently transfected to express hPro-CpepSfGFP without (WT) or with the G(B8)V MIDY mutation. At 48 h post-transfection, the cells were pulse-labeled with ^35^S-labeled amino acids and either lysed immediately, or chased for 90 min as indicated. Cell lysates and media were immunoprecipitated with anti-GFP antibodies and analyzed by **C)** nonreducing, or **D)** reducing SDS-PAGE and phosphorimaging. hPro-G(B8)V-CpepSfGFP was retained intracellularly and did not appear in the medium. **E)** Recovery of newly-synthesized hPro-CpepGFP and hPro-G(B8)V-CpepSfGFP from n = 3 experiments (mean + s.d.).

**Figure 8.**
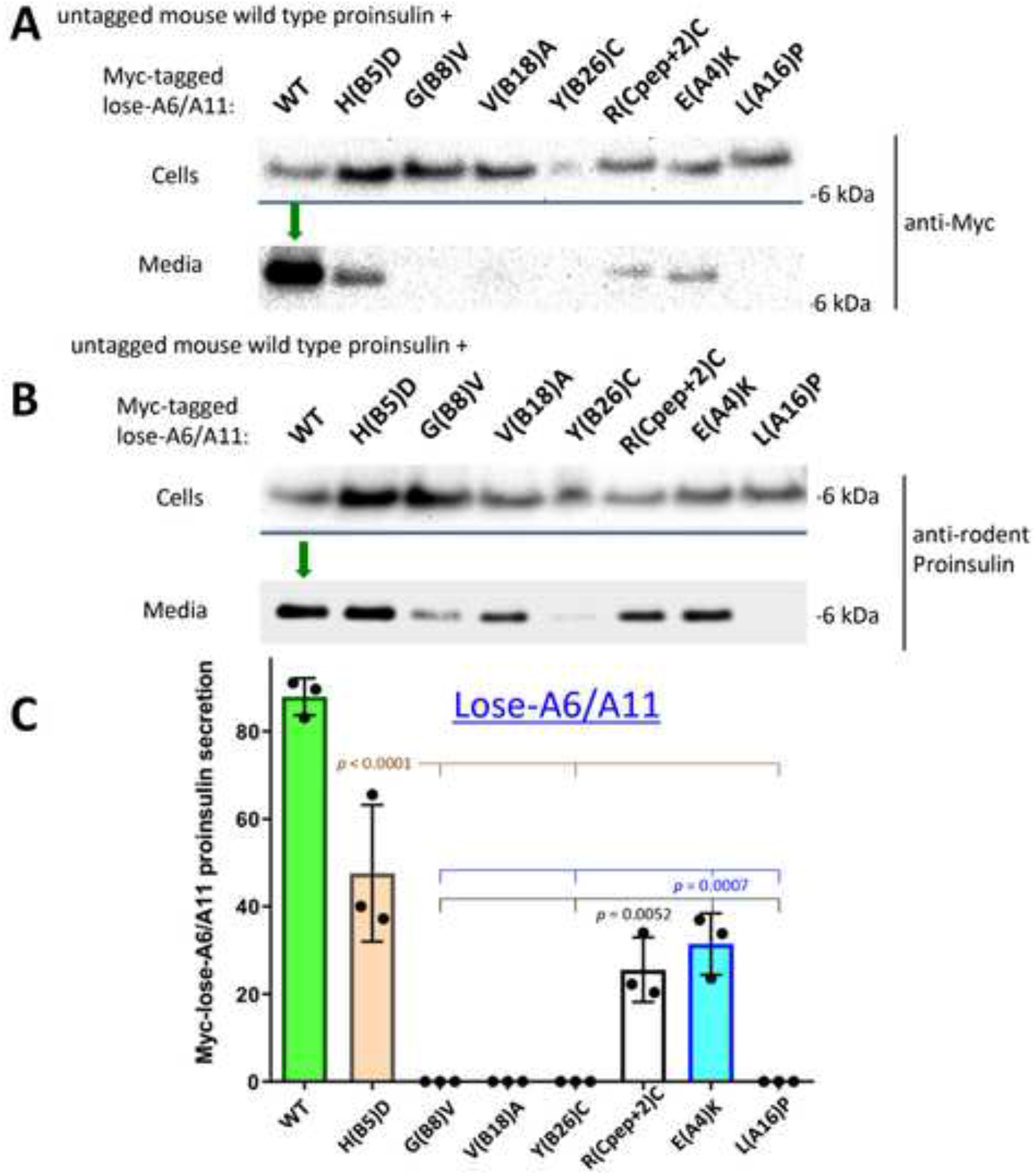
Co-expression of myc-tagged lose-A6/A11-MIDY proinsulin with mouse WT proinsulin does not efficiently block untagged WT proinsulin export. 293T cells were co-transfected to express untagged mouse WT proinsulin-2 and myc-tagged human WT or MIDY mutant in the lose-A6/A11 background. Transfection and sample processing was exactly as in Fig. 2. **A)** Cell and media content of myc-tagged lose-A6/A11 proinsulins, immunoblotted with anti-Myc (as in Fig. 2A); tagged-WT proinsulin was secreted (green arrow). **B)** Cell and media content of co-expressed untagged mouse WT proinsulin, immunoblotted with anti-rodent proinsulin (as in Fig. 2B); untagged-WT proinsulin was secreted (green arrow). **C)** Secretion of WT or MIDY mutant human proinsulins in the lose-A6/A11 background, quantified from three independent replicate experiments conducted similarly to that described in panel A. Mean + s.d. from n = 3 independent experiments. Secretion of H(B5)D, R(Cpep+2), and E(A4)K was significantly increased compared to other MIDY mutants G(B8)V, Y(B26)C, and L(A16)P (*p*-values shown on Figure).

### Structural Modeling

Spatial relationships in native insulin were examined in the wild-type zinc insulin hexamer (Protein Databank [PBD] entry 4INS, from the insulin hexamer) [22, 23]. This structure contains an asymmetric dimer (TT’) in the asymmetric unit. Structural elements were visualized using InsightII and Pymol.

### Data and Resource Availability

Data generated and analyzed during this study are included in the published article (and its online supplementary files). Additional data generated during the current study are available from the corresponding author upon reasonable request.

## RESULTS

### The effect of MIDY mutations on proinsulin folding and export

In this study, we have examined MIDY mutants producing autosomal-dominant neonatal diabetes, including H(B5)D [7], a previously unreported G(B8)V, Y(B26)C and L(A16)P [24], and the commonly reported R89C [which has a polypeptide position R(Cpep+2)C] [25], as well as two mutants producing diabetes at somewhat later ages including V(B18)A [26] and E(A4)K [27]. Most MIDY missense mutants fall within the established primary structure of mature human insulin. Fig. 1A highlights the degree of natural sequence variation at each position [28]; Figs. 1B, C show the site of these MIDY residues in the 2-chain insulin structure. Unsurprisingly, MIDY mutations tend to occur at sites of residue conservation, indicating evolutionary intolerance for substitution. Although the C-peptide and its flanking cleavage sites are not shown, we note that R(Cpep+2)C replaces a highly conserved Arg at the C-peptide/A-chain cleavage site with an extra (7^th^) cysteine that, by definition, has no natural disulfide partner.

Each MIDY mutant was engineered into the hPro-CpepMyc cDNA encoding human proinsulin bearing myc-tagged C-peptide [12]. When expressed heterologously in 293T cells, the construct bearing the WT insulin chains was secreted to the medium, in contrast with any of the 7 MIDY mutants (Fig. 2A; quantified in Fig. 2C). Moreover, WT human proinsulin expression allowed for secretion of co-expressed WT mouse proinsulin (detected with a species-specific proinsulin mAb), whereas each of the MIDY mutants yielded a “bystander effect” [11, 29], inhibiting the secretion of co-expressed WT proinsulin (Fig. 2B).

The abundance of expression of WT proinsulin, or other factors ongoing in β-cells, can subvert the efficiency of ER quality control, leading to escape of a subset misfolded proinsulin molecules from the ER [14]. With this in mind, we expressed MIDY proinsulins in Min6 (mouse) β-cells — using the myc-tag to unequivocally distinguish transfected from endogenous proinsulin. WT human proinsulin was abundantly exported in Min6 cells, being secreted as unprocessed proinsulin (Fig. 3A) and being converted intracellularly to human insulin (Fig. 3B). Whereas all MIDY mutants were significantly impaired in export (Fig. 3C), there was demonstrable low-level secretion of several MIDY mutants, notably including proinsulin-V(B18)A (Fig. 3A, C). We could independently confirm some intracellular transport of proinsulin-V(B18)A by detection of processed human insulin in these mouse β-cells (Supplemental Fig. S1), indicating that V(B18)A was in small part able to reach immature secretory granules for proinsulin-to-insulin conversion [30, 31].

Using nonreducing Tris-tricine-urea-SDS-PAGE to discern the formation of native B-A-chain disulfide bonds within newly-synthesized proinsulin [19], we examined the oxidation of myc-tagged human proinsulins expressed in INS832/13 β-cells. Whereas newly-synthesized WT proinsulin predominantly formed native disulfide bonds, this was impaired in 6 of the 7 MIDY mutants (Fig. 4A, B). Newly-synthesized MIDY mutants were, once again, significantly impaired in their secretion from β-cells (Fig. 4C, D). Interestingly, proinsulin-R(Cpep+2)C exhibited native-like oxidation [Figs. 4A; bearing in mind an extra (7^th^) cysteine exposed on the proinsulin surface, based upon its location at the cleavage site]; nevertheless it exhibited severely defective secretion (Fig. 4C, D).

### MIDY proinsulins engage in intermolecular disulfide mispairing in the ER of β-cells

We recently found that proinsulin in pancreatic β-cells is susceptible to forming intermolecular disulfide-linked complexes, and an increased abundance of these aberrant complexes precedes the onset of diabetes [32]. In lysates of Min6 cells transfected to express myc-tagged human proinsulin or MIDY mutants, we looked for the presence of proinsulin disulfide-linked complexes (resolved by nonreducing SDS-PAGE and detected by human-specific proinsulin immunoblotting). Reducing SDS-PAGE (in which intermolecular disulfide-linked proinsulin runs as reduced monomers) was used to detect total human proinsulin levels in the cells. By nonreducing SDS-PAGE, whereas WT myc-tagged human proinsulin was predominantly monomeric, the various MIDY mutants formed an increased fraction of disulfide-linked dimers and higher molecular weight complexes (Fig. 5A). By analysis using two different proinsulin antibodies, we were able to calculate a misfolding index indicating increased disulfide-linked complex formation for at least three (and probably all) of the MIDY proinsulin mutants (Supplemental Fig. S2). Moreover, there was a suggestion of increased ER stress induced by most MIDY mutants [response measured by BiP-luciferase assay, although only L(A16)P generated statistical significance (Supplemental Fig. S3)].

### Impact of cysteines A6 and A11 on MIDY proinsulin folding and export

A proinsulin double mutant known as lose-A6/A11 (bearing Cys-to-Ser substitutions) exhibits essentially normal intracellular trafficking in β-cells; thus, MIDY mutants retained by virtue of an unpaired Cys(A11) [19] might be rescued in a lose-A6/A11 background. We immediately observed significantly rescued secretion of H(B5)D, R(Cpep+2)C, and E(A4)K [and a trend of increased secretion for V(B18)A] (Fig. 6A, B) in Min6 β-cells, but no important benefit to other MIDY mutants G(B8)V, L(A16)P, and not significant effect for Y(B26)C (Fig. 6A, B). Nevertheless, most newly-synthesized MIDY mutants now exhibited a trend of an increased fraction of near-native oxidized forms (Fig. 6C; compare to Fig. 4A), and this was statistically significant for two of the mutants that exhibited secretion rescue [H(B5)D and E(A4)K (Supplemental Fig. S4)]. However, despite this indication of improved oxidation in β-cells — on repeated analysis, while improvement of export of newly-synthesized MIDY mutants in the lose-A6/A11 background was observed for H(B5)D, R(Cpep+2)C, E(A4)K [and a trend to increased secretion for V(B18)A], there was no significantly enhanced export for G(B8)V, Y(B26)C, or L(A16)P (Fig. 6D, E). [Identical results were obtained upon expression of these constructs in 293T cells, *not shown*.]

Interestingly, whereas lose-A6/A11 improved oxidation of most of the MIDY mutants as noted above, MIDY proinsulin mutants G(B8)V, Y(B26)C, or L(A16)P — recovered well by reducing SDS-PAGE — were caught up primarily in intermolecular disulfide-linked complexes rather than proinsulin monomers (Fig. 5B). Remarkably however, the lose-A6/A11 background appreciably rescued detectable intracellular proinsulin monomers for H(B5)D, R(Cpep+2), and E(A4)K, as well as V(B18)A (Fig. 5B). These data strongly suggest that these latter MIDY mutations inhibit the pairing of Cys(A6) and Cys(A11), leading to their engagement in inappropriate intermolecular disulfide bond formation (which is improved in the lose-A6/A11 background).

When expressed either in pancreatic β-cell lines or in β-cells *in vivo*, a construct known as hPro-CpepSfGFP (bearing a superfolder-GFP within the C-peptide of human proinsulin) tends to accumulate in the juxtanuclear Golgi region and becomes stored as processed fluorescent C-peptide in mature secretory granules [13, 33]. With this in mind, we examined each of the previously noted MIDY mutants in an otherwise WT or lose-A6/A11 proinsulin background in INS832/13 β-cells. In the absence of mutations causing MIDY, both proinsulin and lose-A6/A11 were delivered to punctate secretory granules at the β-cell periphery; and each of the MIDY mutations introduced into the otherwise WT proinsulin background exhibited a cytoplasmic reticular distribution classic for the ER (Fig. 7A and Supplemental Fig. S5). In conjunction with this, WT proinsulin could be seen to be cleaved (albeit weakly) to a conversion intermediate and fully processed SfGFP-tagged C-peptide, and this was not obvious for the MIDY mutants (Fig. 7B *upper panel, green arrow*). When introduced into the lose-A6/A11 background, the MIDY mutations Y(B26)C, G(B8)V, and L(A16)P continued to exhibit a classic ER-like distribution, whereas H(B5)D, R(Cpep+2)C, and E(A4)K each reverted to a granule pattern (Fig. 7A and Supplemental Fig. S5). In conjunction with this, the latter three MIDY mutants also yielded processed GFP-tagged C-peptide (Fig. 7B *lower panel, green arrows*). [As an aside, the common R(Cpep+2)C mutation mutates the C-A junctional cleavage site, and the cleavage intermediate band generated from this construct exhibited a unique, abnormal mobility, suggesting an atypical cleavage (Fig. 7B).] Additionally, the V(B18)A mutation yielded a hint of processed C-peptide in both the WT and lose-A6/A11 background (Fig. 7B) and in counting multiple transfected cell images, some green fluorescent secretory granules were detected, albeit decreased by > 70%. Taken together, these morphological and biochemical findings support the characterization of two main subgroups of MIDY mutants that can either be rescued or not-significantly-benefitted by the lose-A6/A11 substitutions, respectively, while V(B18)A behaves as a slightly better tolerated MIDY mutation.

As the G(B8)V mutation had not been described previously (we learned of this spontaneous mutation in a New Zealand girl who presented with diabetes at 6 weeks of age), we directly compared newly-synthesized G(B8)V with G(B8)S in the lose-A6/A11 background when these constructs were expressed in 293T cells. Whereas the lose-A6/A11 parent protein rapidly formed a near-native oxidized form and underwent significant secretion within an hour after synthesis, both G(B8)V and G(B8)S appeared identical, with impaired oxidation and no detectable secretion to the medium (Supplemental Fig. S6). We further examined the G(B8)V mutation in the hPro-CpepSfGFP background by pulse-chase radiolabeling of the transfected cells, followed by immunoprecipitation with anti-GFP. Immediately post-pulse, recovery of newly-synthesized G(B8)V mutant was less than from cells expressing WT hPro-CpepSfGFP, and both constructs could be detected in disulfide-linked complexes (Fig. 7C, D). By 90 min of chase, however, WT hPro-CpepSfGFP could be recovered in the medium while its intracellular level had fallen (Fig. 7D, E). In contrast for the G(B8)V mutant at 90 min of chase, there was no secretion and its intracellular level had not declined; in fact its intracellular level was now > that of WT hPro-CpepSfGFP (Fig. 7D, E), although almost none of the newly-synthesized mutant protein could be recovered as proinsulin monomers by nonreducing SDS-PAGE (Fig. 7C) – indicative of a severe MIDY mutation.

### Impact of cysteines A6 and A11 on dominant-negative MIDY behavior

To test dominant-negative blockade of WT proinsulin, we co-expressed in 293T cells each human myc-tagged MIDY mutant in the lose-A6/A11 background, with untagged WT mouse proinsulin. We could then look independently by Western blotting at the export of the lose-A6/A11 MIDY constructs (anti-myc immunoblotting) and WT proinsulin (mAb anti-rodent proinsulin). We observed negligible secretion of G(B8)V, Y(B26)C, and L(A16)P in the lose-A6/A11 background (Fig. 8A; quantified in Fig. 8C), accompanied by suppressed escape of co-expressed WT proinsulin (Fig. 8B). In contrast, we detected significant secretion of H(B5)D, R(Cpep+2)C, and E(A4)K in the lose-A6/A11 background (Fig. 8A; quantified in Fig. 8C) accompanied by secretion of co-expressed WT proinsulin (Fig. 8B). These data suggest that the same proinsulin mutants that are themselves less susceptible to ER quality control are also less effective in *trans*-dominant blockade of co-expressed WT proinsulin. Indeed, when comparing to Figure 2, none of the lose-A6/A11 MIDY constructs except for L(A16)P are as efficient in blocking co-expressed WT proinsulin, including V(B18)A which itself appears blocked yet cannot efficiently block the trafficking of co-expressed WT proinsulin. Thus the data highlight the potential importance of unpaired Cys(A11) and Cys(A6) in adverse proinsulin folding and trafficking.

## DISCUSSION

MIDY is disease of protein misfolding causing autosomal dominant diabetes. MIDY mutations threaten native proinsulin disulfide bond formation [12, 19]. ER quality control recognizes and prevents export of misfolded proinsulin, thus MIDY mutants have loss of function in generating insulin [4, 5]. Moreover, natural self-association properties of proinsulin enable the defective MIDY gene product to associate with and dominantly block export of WT proinsulin [10, 11, 34]. Nevertheless, different MIDY mutations lead to different ages of diabetes onset, suggesting greater or lesser severity of proinsulin misfolding [4, 35]. In this study, we’ve examined seven MIDY proinsulin mutants by testing for the importance of Cys(A6) and Cys(A11) because, curiously, these residues normally engage in an intramolecular disulfide bond that is nonessential for proinsulin trafficking [18, 19] yet in MIDY they may participate in intermolecular disulfide bonds that could render mutant proinsulin molecules unacceptable to ER quality control.

### MIDY mutants exhibit a range of trafficking defects

Proinsulin-V(B18)A is linked to diabetes onset beyond the neonatal period [ages 3.3 – 10.5 years [26]]. Although interchange of Val-to-Ala is a conservative substitution that is generally well-tolerated [36], protein context is important in determining pathogenicity, as some similar missense substitutions are linked to disease [37]. Equally important is cellular context: we did not detect anterograde trafficking of proinsulin-V(B18)A from 293T cells even in the lose-A6/A11 background (Fig. 8), yet low-level anterograde trafficking was observable in pancreatic β-cells (Fig. 3), even leading to the formation of small quantities of mature human insulin (Supplemental Fig. S1). These data suggest that the V(B18)A mutant exhibits a slightly less severe proinsulin misfolding than a number of other MIDY mutants.

### Defective Cys(A6)-Cys(A11) pairing defines two subgroups of mutations

The MIDY mutants G(B8)V, Y(B26)C, and L(A16)P exhibited significantly less anterograde trafficking than WT proinsulin, and even less than the MIDY mutant proinsulin-V(B18)A. The trafficking defect of the newly-described proinsulin-G(B8)V is severe and very similar to that of the G(B8)S mutation (Supplemental Fig. S6), which has been recently described to account for spontaneous diabetes in the KINGS diabetic mouse [38], as well as being one of the first described human MIDY mutants [25] — its folding defect is considered further, below. Additionally, it might be predicted that proinsulin-Y(B26)C and L(A16)P (also discussed further, below) would be catastrophically detrimental to folding as they (in addition to position B8) fall among the more conserved residues within insulin (Fig. 1A) — and patients bearing these heterozygous mutations have been found to be insulin-requiring within the first 6 months of life [24]. In 293T cells, these mutants could not be secreted either in the proinsulin or lose-A6/A11 background (Figs. 2, 8); moreover, secretion was profoundly low in pancreatic β-cells (Fig. 3), and could not be statistically increased in the lose-A6/A11 background in either Min6 or INS832/13 cells (Figs. 6, 8).

In contrast with the foregoing proinsulin mutants, we find that MIDY mutations encoding H(B5)D, R(Cpep+2)C, or E(A4)K fall into a distinct group. Specifically, whereas these mutants cannot be secreted in the proinsulin backgound, they are all exported to a greater degree in the lose-A6/A11 background — able to be secreted (Fig. 6) and to be processed to mature C-peptide in pancreatic β-cells (Fig. 7).

Recovery of export in the lose-A6/A11 background can be accounted for by diminished formation of intermolecular disulfide-linked complexes (Fig. 5). As this particular subgroup of MIDY mutants develops improved folding in the lose-A6/A11 background, leading to improved trafficking in β-cells, we conclude that for this subset of MIDY proinsulin mutants, burying these cysteines in the Cys(A6)-Cys(A11) disulfide bond makes them unavailable for intermolecular disulfide mispairing [19]. Rescue in the lose-A6/A11 background then allows unperturbed initial formation of the Cys(B19)-Cys(A20) disulfide, leaving driving the remaining Cys(B7) and Cys(A7) free thiol groups into disulfide partnership [19], eliminating or diminishing the predisposition for intermolecular disulfide bond formation, and that is one of the major factors in ER quality control of proinsulin export.

In the ‘T-state’ insulin crystal structure [39], the imidazolic ring of H(B5) packs within an interchain near to the Cys(A6)-Cys(A11) disulfide bond (Supplemental Fig. S7A). We posit that H(B5)D may alter the relative positions and solvation of Cys(A6) and Cys(A11) [and to some extent also Cys(A7) [40, 41]]. Additionally, in the solution structure of proinsulin, R(Cpep+2) and E(A4) appear to form a salt bridge and may contribute to the ‘CA knuckle’ [42], which may also help to stabilize the proximal A-chain α-helix [43]. In turn, there is a close relationship between proximal A-chain conformation and the Cys(A6)-Cys(A11) disulfide bond [44]. (Additionally it seems likely that the extra unpaired cysteine of R(Cpep+2)C might also engage in intermolecular disulfide crosslinking — for which further study is still needed.) Thus, MIDY mutations H(B5)D, R(Cpep+2)C, and E(A4)K are each expected to perturb the Cys(A6)-Cys(A11) disulfide bond, impacting on ER quality control of proinsulin export, and this may help to explain the beneficial effects of the lose-A6/A11 substitutions.

In contrast, a second subset of MIDY mutants [G(B8)V, Y(B26)C, and L(A16)P] cannot be rescued in the lose-A6/A11 background. G(B8), invariant among vertebrate insulins, insulin-like growth factors (IGFs) and relaxins, lacks a side chain but exhibits a critical positive ϕ dihedral angle ordinarily forbidden to L-amino acids [23]. From *in vitro* folding studies, any L-amino-acid substitution in this position predisposes to impaired disulfide pairing [45] as observed for the G(B8)S MIDY mutant [46], triggering permanent neonatal diabetes [25, 38]. We posit that G(B8)V similarly enforces an inappropriate negative B8 ϕ dihedral angle, in turn misorienting the side chain of Cys(B7), which cannot be rescued by the lose-A6/A11 template. The side chain of Y(B26) is inserted into a solvent-exposed inter-chain crevice (Supplemental Fig. S7B), and its packing against the conserved nonpolar surface of the central-B-chain α-helix bearing L(B11), V(B12) and L(B15) stabilizes a nascent native-like supersecondary structural “U-turn” [23] to promote proper formation of the essential Cys(B19)-Cys(A20) disulfide bond [9, 47], which is likely to be disrupted in the Y(B26)C MIDY mutant — and this too cannot be rescued in the lose-A6/A11 background. L(A16) fits snuggly within a potential internal cavity at the confluence of insulin’s three native α-helices (Supplemental Fig. S7C), and the A16-A19 α-helical turn is thought to facilitate alignment with the B-chain α-helix to enable initial Cys(B19)-Cys(A20) pairing [47]. The imino-side chain of P(A16) is expected to distort local α-helical conformation [48], incurring a local steric clash and destabilizing the internal cavity, leading to an insuperable block to formation of the Cys(B19)-Cys(A20) pairing within proinsulin’s folding nucleus [9].

In a slightly milder category from the others, V(B18)A would create a packing defect (Supplemental Fig. S7D) that would be expected to enlarge the range of possible positions/orientations accessible to the adjoining Cys(B19) and its Cys(A20) partner. Evidently, this mutation does not completely preclude Cys(B19)-Cys(A20) pairing (Fig. 3C, Supplemental Fig. S1), with slight secretory improvement by the lose-A6/A11 template (Fig. 6A-C; Supplemental Fig. S5).

Finally, we note that the lose-A6/A11 background resulted in notable diminution of dominant-negative behavior, particularly for the subgroup of MIDY mutants that are themselves rescued in folding and intracellular transport [a modest effect was also observed for Val(B18)A]. These data strongly support the hypothesis that it is the recognition and retention of the misfolded mutant partner by ER quality control that confers ER retention upon the co-expressed WT partner [5]. Indeed there is a definite proinsulin-proinsulin partnership, as indicated by the importance of proinsulin dimerization propensity to dominant-negative behavior [34]. Altogether, the findings in this report highlight an improved structural understanding by which proinsulin misfolding might be attacked pharmacologically, a) to block exposure of unpaired cysteines, and b) to limit the increased aberrant proinsulin-proinsulin associations [49] which are a hallmark of MIDY and perhaps also of type 2 diabetes [32].

## Declarations

### Ethics approval and consent to participate

University of Michigan biosafety approval for recombinant DNA: IBCA00001142

### Consent for publication

All co-authors consent

### Availability of data and material

All primary data and material in the manuscript are available upon reasonable request.

### Authors’ contributions

L.H, A.A., A.H, and K.R. generated research data. P.A. wrote the manuscript. All authors reviewed the manuscript. M.L., B.D., M.A.W., B.T., L.H., A.A. and P.A. revised/edited the manuscript and contributed to discussion.

## Acknowledgements

The authors thank the University of Michigan Protein Folding Diseases Initiative, and Michigan Diabetes Research Center Imaging core (NIH P30-DK-020572) for assistance with live cell imaging. We thank D. Chatterjee and J. Racca for assistance with molecular graphics.

## Abbreviations

ER: endoplasmic reticulum
MIDY: Mutant *INS*-gene induced Diabetes of Youth

## Key Resources Table

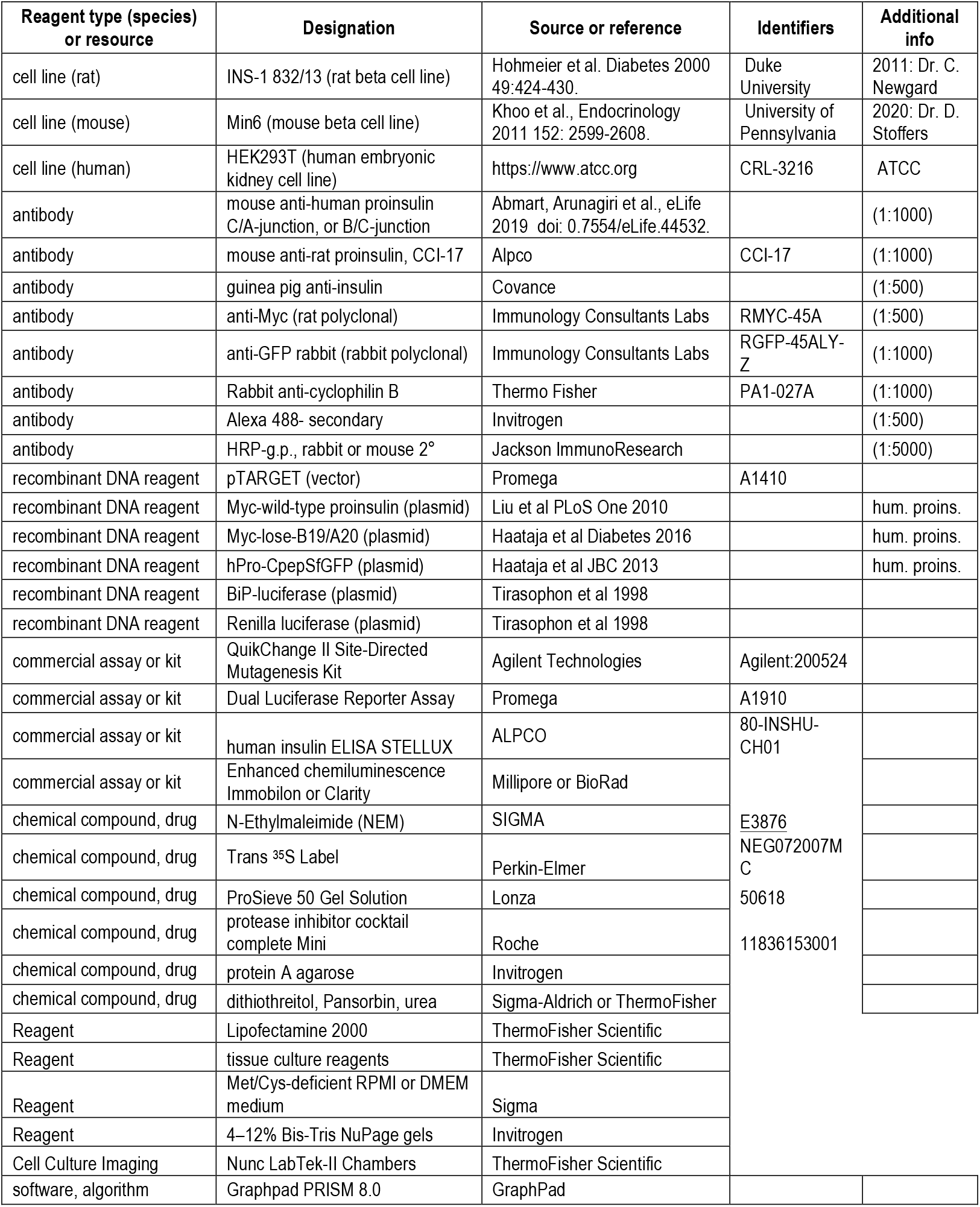

