## Supplement for "Distinct states of proinsulin misfolding in MIDY"

### Supplemental Figures

**Supplemental Figure Legends**

**Supplement Figure S1.** MIDY mutant V(B18)A retains some ability to produce and release human insulin in β-cells. Recombinant myc-tagged human proinsulin WT or MIDY mutants were transiently expressed in Min6 (mouse) β-cells. At 24 h after transfection, the bathing medium was changed to fresh medium that was then collected for 24 h without additional stimulation, before cell lysis and analysis of both lysates (yellow bars) and media (blue bars) by human-specific insulin ELISA. Mean + s.d. from n = 3 independent experiments; * $p < 0.05$ for secreted human insulin derived from proinsulin-V(B18)A compared to that from other MIDY mutants.

**Supplement Figure S2.** MIDY mutant proinsulins are predisposed to form disulfide-linked complexes. Min6 (mouse) β-cells cells were transiently with human WT or MIDY mutant proinsulins and resolved by nonreducing and reducing SDS-PAGE, as in Fig. 5. Disulfide-linked dimer + trimer bands detected by immunoblotting of the nonreducing gels with anti-Myc or anti-human proinsulin were divided by the total proinsulin band intensity detected by immunoblotting on the reducing gels, and the predisposition to form disulfide-linked complexes was compared to that of WT proinsulin (first bar); error bars (s.d.) and significant *p*-values are indicated on the figure.

**Supplement Figure S3.** ER stress response measurement in β-cells expressing MIDY mutants. INS832/13 β-cells were co-transfected with myc-tagged human WT or MIDY mutant proinsulins plus BiP-firefly luciferase and CMV-renilla luciferase. At 48 h post-transfection, luciferase activities in transfected cells were measured as described in Methods (mean ± s.d.). As shown, all MIDY mutants showed a trend to increased ER stress, but only L(A16)P achieved statistical significance (*p*-value indicated on the figure) compared to WT proinsulin.

**Supplement Figure S4.** Oxidation of newly-synthesized proinsulin and lose-A6/A11 proinsulin. INS832/13 β-cells were transfected to express myc-tagged human WT or MIDY mutant proinsulins (P), as well as WT or MIDY versions of lose-A6/A11 proinsulins (L). Pulse-labeling (without chase) and sample processing was performed as described in Figs. 4A and 6C, and the fastest-migrating (oxidized) forms were quantified (mean ± s.d.) from at least 3 independent experiments. In the lose-A6/A11 background, the MIDY mutants H(B5)D and E(A4)K both exhibited significantly improved oxidative folding (*p*-values indicated on the figure). The third MIDY mutant whose secretion is increased in the lose-A6/A11 background, R(Cpep+2)C, already showed native-like disulfide bond formation even in the WT proinsulin background.

**Supplement Figure S5.** Live cell imaging of INS832/13 cells transiently expressing fluorescent WT or MIDY proinsulins (and their derived fluorescent protein products). The NIS Elements software "volume view" was used to create a 3D-tilt model to highlight the intracellular distribution of fluorescent protein. Top eight panels: WT or MIDY versions of hPro-CpepSfGFP (bearing superfolder-GFP embedded within the C-peptide). Only the WT construct displays a punctate fluorescence pattern typical of secretory granules in live β-cells (red box). Bottom eight panels: the same WT and MIDY hPro-CpepSfGFP constructs in the lose-A6/A11 background. In addition to WT, in this background E(A4)K, H(B5)D and R(Cpep+2)C all show a punctate fluorescence pattern typical of secretory granules in live β-cells (red boxes). The other constructs continued to show a fluorescence pattern typical of ER retention. The V(B18)A substitution was an exception showing a partial granule-like pattern, especially in the lose-A6/A11 background (yellow box). White scale bar = 10 μm.

**Supplement Figure S6.** Oxidation and secretion of newly-synthesized lose-A6/A11 proinsulin bearing G(B8V) and G(B8)S MIDY substitutions. 293T cells transfected to express the indicated constructs were pulse-labeled with $^{35}$S-labeled amino acids, and both cell lysates and media were immunoprecipitated with guinea pig polyclonal anti-insulin at the indicated chase times. The samples were analyzed by Tris-tricine-urea-SDS-PAGE under nonreducing and reducing conditions, followed by autoradiography.

**Supplement Figure S7.** Structural environments of mutation sites in T-state insulin monomer (coordinatesfrom Protein Databank entry 4INS; 2-Zn molecule-2). **A)** The imidazole side chain of H(B5) (blue dashed circle) packs within an inter-chain crevice near both the internal Cys(A6)-Cys(A11) disulfide bridge (these positions circled in the image) [and near Cys(A7) and Ser(A9) with which it forms bifurcating hydrogen bonds (dashed yellow lines)]. **B)** Packing of the side chain of Y(B26) within a shallow solvent-exposed inter-chain crevice lined by the side chains of F(B24), P(B28), G(A1) and I(A2). **C)** Internal side chain of L(A16) is shown in relation to insulin's three α-helices (ribbons, and a partially transparent protein surface). Core packing of L(A16) involves Cys(A6)-Cys(A11) (shown as yellow balls) and the side chains of L(B11), L(B15), I(A2), L(A13), and Y(A19); Cys(A20)-Cys(B19) (not shown) is also nearby. **D)** Packing of the side chain of V(B18) within a deep solvent-exposed interchain crevice lined by A(B14) and L(A13) at its lip and in its depths by the core side chains of L(A16) and Cys(A20).

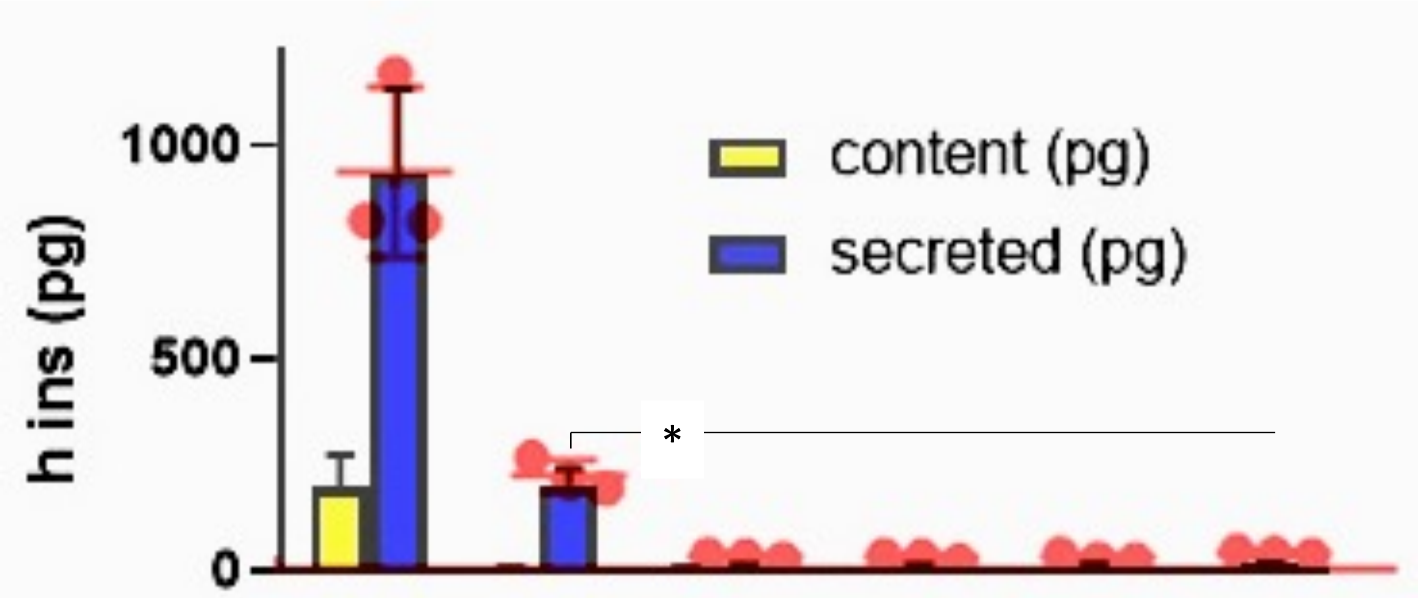


**Myc-tagged hProinsulin: WT V(B18)A Y(B26)C R(Cpep+2)C E(A4)K L(A16)P**

Haataja, Arunagiri et al., Supplemental Fig. S1

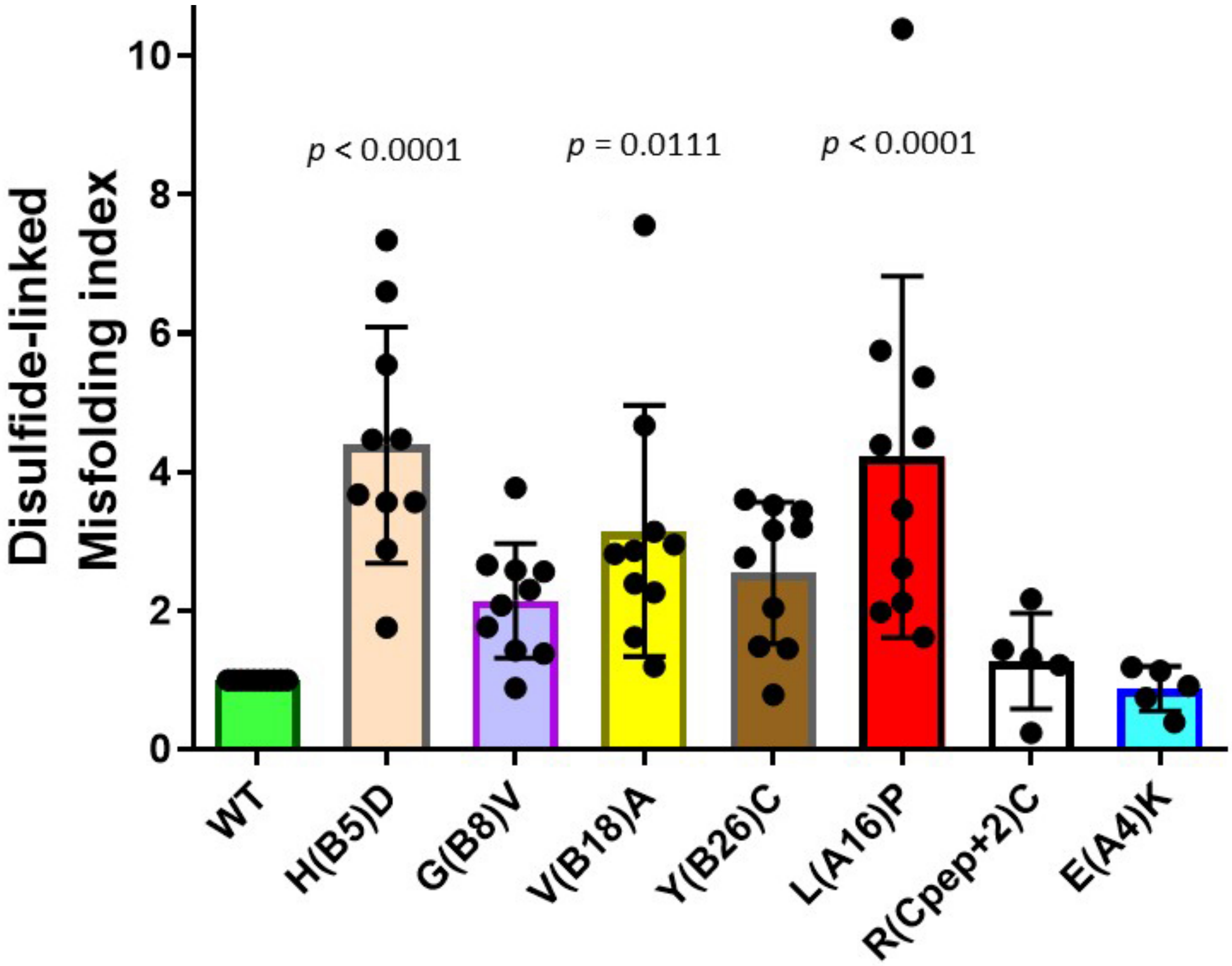


Haataja, Arunagiri et al., Supplemental Fig. S2

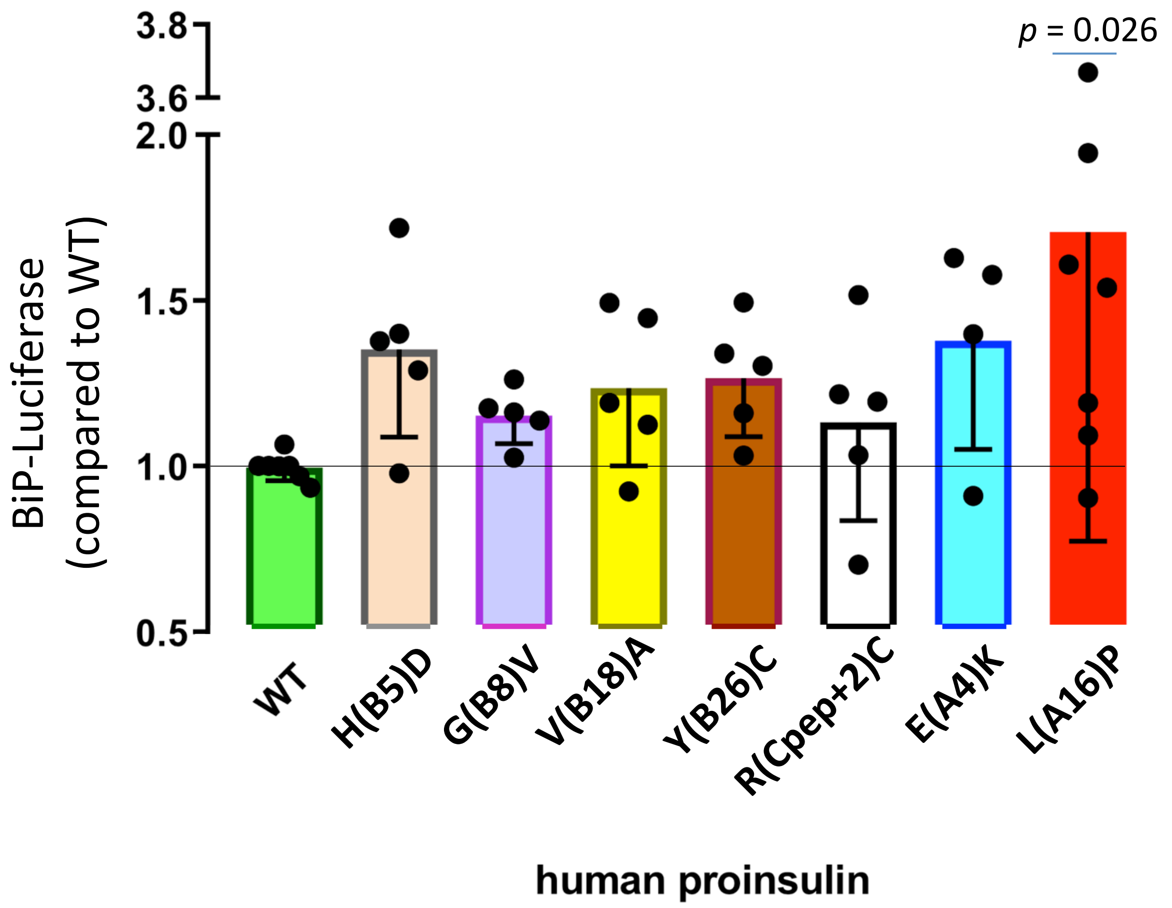


Haataja, Arunagiri et al., Supplemental Fig. S3

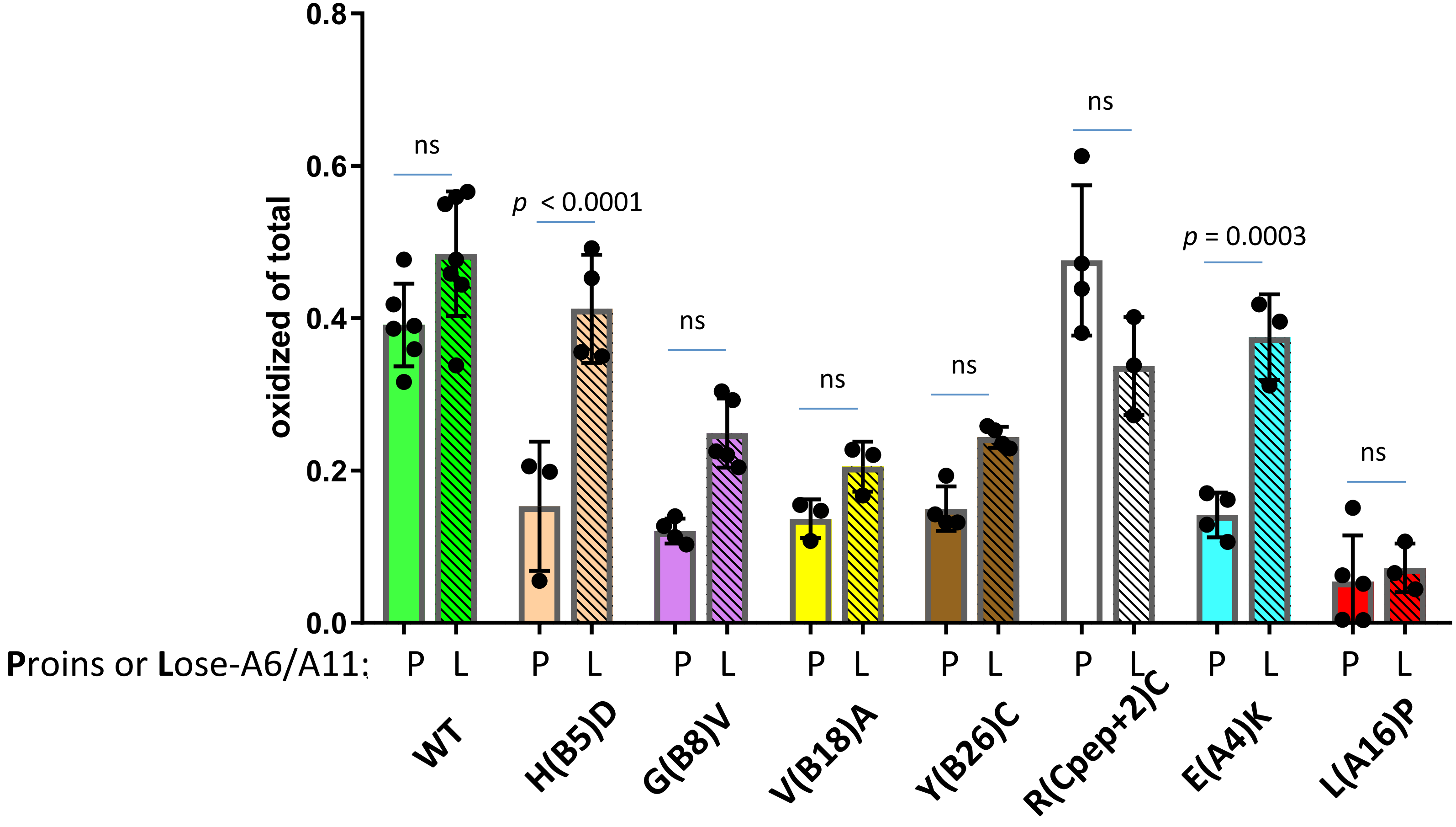


Haataja, Arunagiri et al., Supplemental Fig. S4

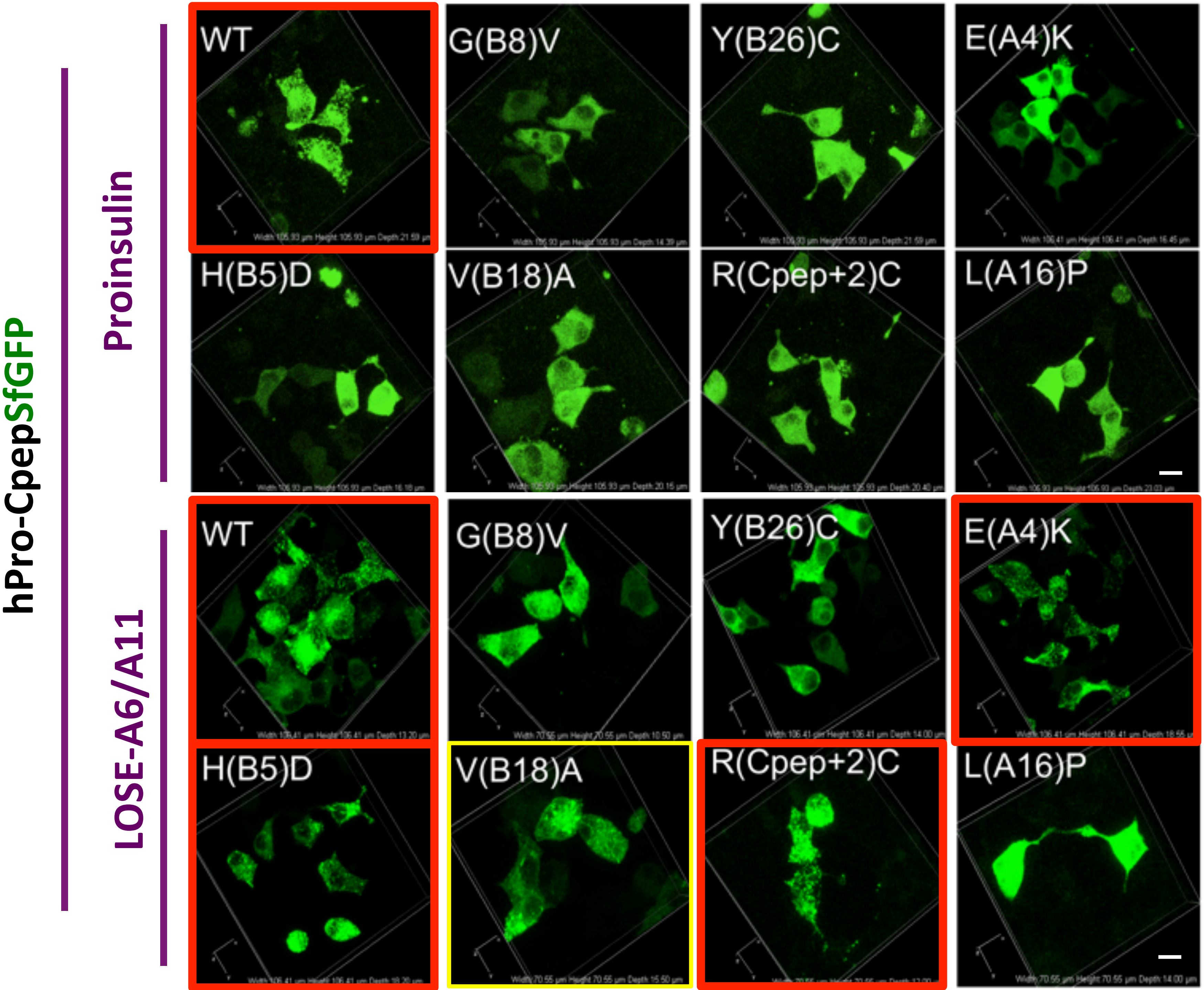


Haataja, Arunagiri et al., Supplemental Fig. S5

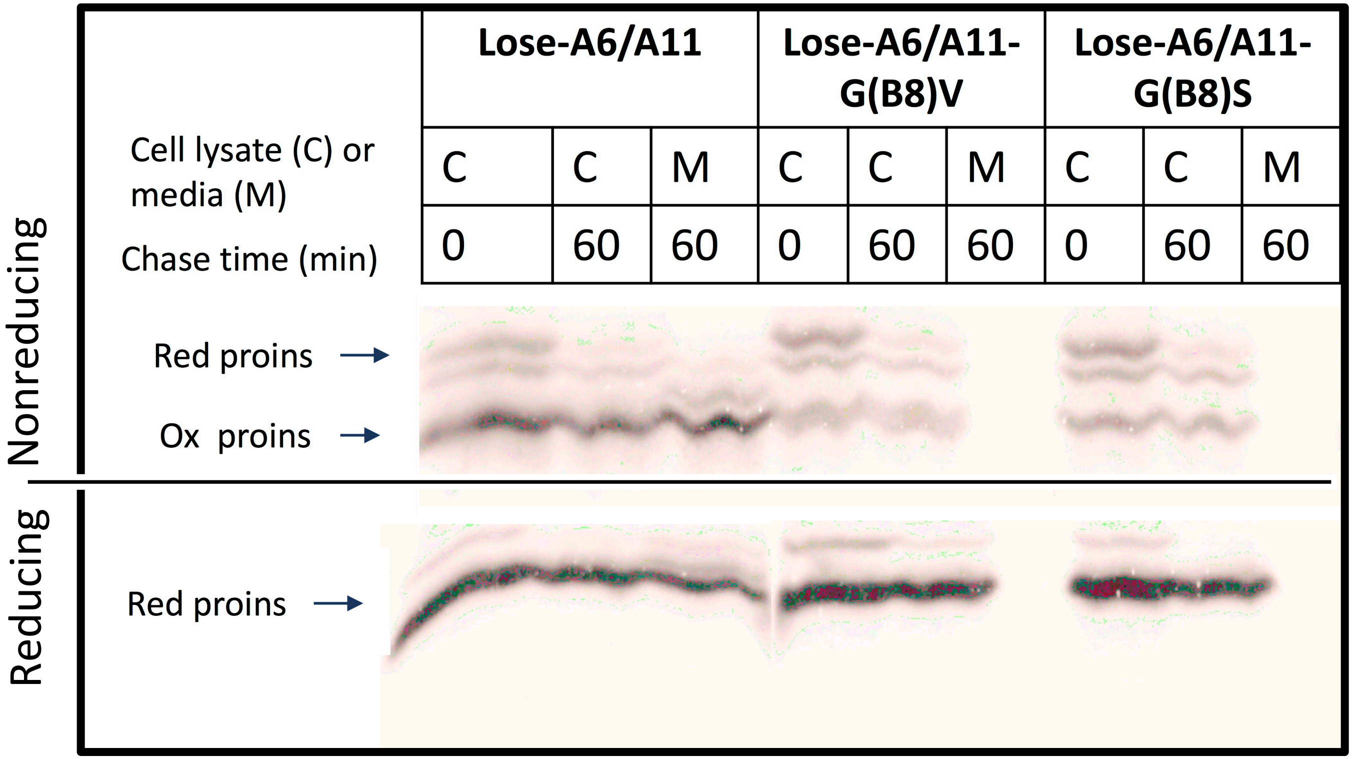


Haataja, Arunagiri et al., Supplemental Fig. S6

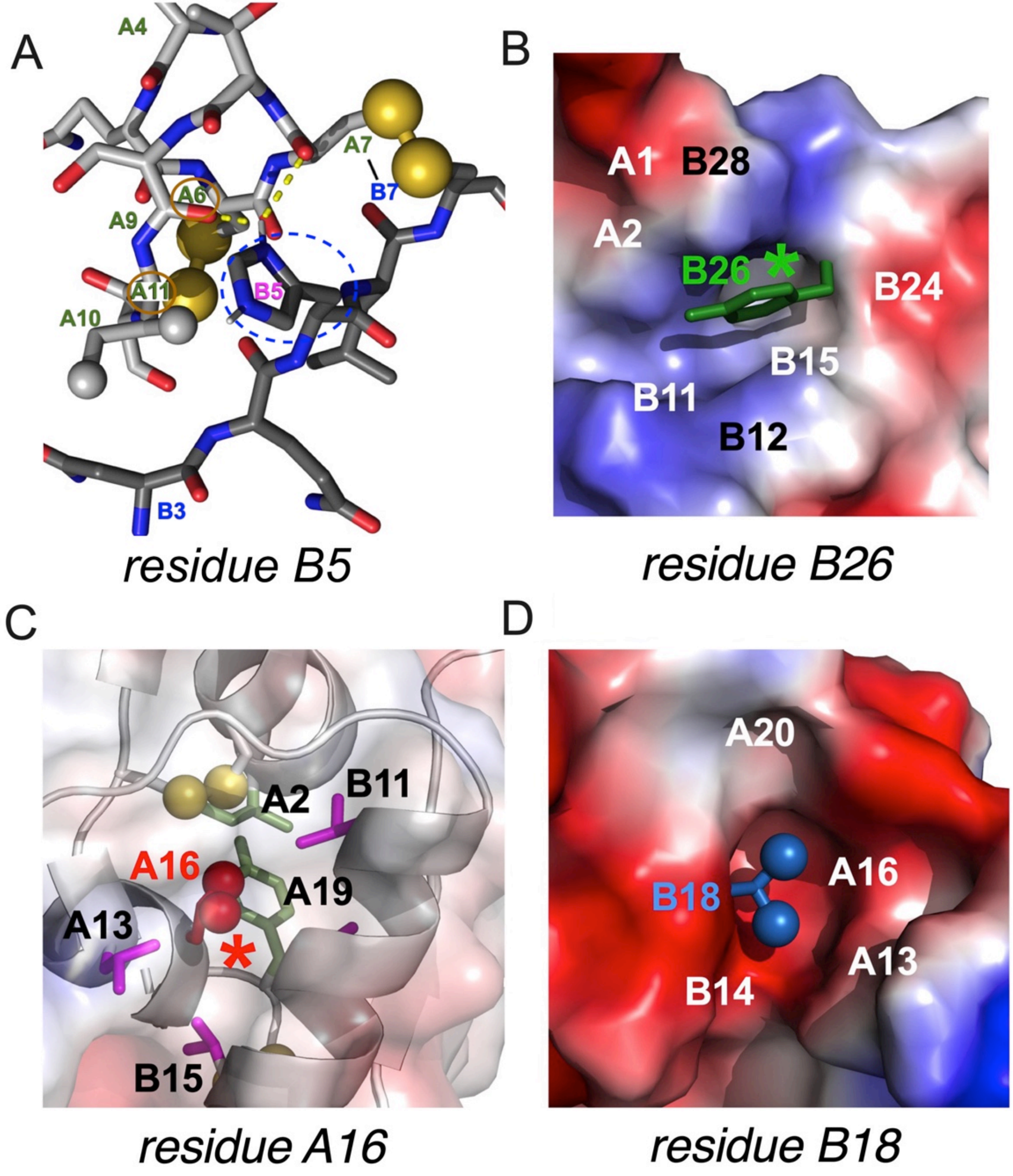


Haataja, Arunagiri et al., Supplemental Fig. S7
